# Complete depletion of *Arabidopsis* linker histones impairs the correlations among chromatin compartmentalization, DNA methylation and gene expression

**DOI:** 10.1101/2021.07.08.451606

**Authors:** Zhenfei Sun, Min Li, Yunlong Wang, Hui Zhang, Yu Zhang, Min Ma, Pan Wang, Yaping Fang, Guoliang Li, Yuda Fang

## Abstract

In eukaryotic cells, linker histone H1 anchors in and out ends of nucleosome DNA to promote chromatin to fold into the 30 nm fiber. However, if H1 plays a role in coordinating the three-dimensional (3D) chromatin architecture, DNA methylation, and transcriptional regulation is not clear. We engineered H1 knockout mutants in *Arabidopsis thaliana* which shows pleiotropic phenotypes. Using High-throughput Chromosome Conformation Capture (Hi-C), we found that H1 complete depletion dampens inter- and intra-chromosomal interactions, as well as intra- and inter-chromosomal arm interactions. MNase accessibility assays followed by sequencing (MNase-seq) showed that the nucleosome density decreases in centromeric regions and increases in chromosome arms. In contrast, DNA methylation level in CHG and CHH contexts increases in centromeric regions and decreases in chromosome arms as revealed by whole genome bisulfite sequencing (WGBS) in *h1* mutant. Importantly, the functional link between DNA methylation and gene transcription is defected, and the extensive switches between chromatin compartment A and B are uncoupled from genome-wide DNA methylation and most of gene transcriptions upon H1 depletion. These results suggested that linker histone H1 works as linkers among chromatin compartmentalization, DNA methylation and transcription.

---

Genomic DNA in eukaryotic cells is densely packaged into chromatin to fit within the small volume of nucleus. The eukaryotes achieve DNA package depending on two types of major proteins, the core histones (H2A, H2B, H3 and H4) and the linker histone H1. Two copies of each of the four core histones wrapped with ∼147 bp of DNA termed as nucleosome core particle together with an additional variable length of linker DNA constitute nucleosome, which is the repeating structure unit of chromatin (Kornberg 1974; Luger et al. 1997). In addition, H1 binds to nucleosome with ∼10 bp of DNA at both the entry and exit sites of nucleosome core particle to facilitate the folding of chromatin into a 30 nm fiber (Bednar et al. 1998; Song et al. 2014; Thoma et al. 1979). H1 is the most variable histone, with multiple H1 family members are present in different organisms. For example, mice and human contain H1-coding genes up to eleven, consisting of seven somatic subtypes and four germ line-specific subtypes. Deletion of any one or two H1 subtypes did not noticeably affect mouse development due to the compensatory effect of other subtypes (Drabent et al. 2000; Fan et al. 2001; Fantz et al. 2001; Lin et al. 2000; Sirotkin et al. 1995). Elimination of H1c, H1d and H1e led to a decrease in the ratio of H1 to nucleosome core particle up to 50 % and embryonic lethality. In addition, deletion of the three H1 subtypes induced the decrease of global nucleosome spacing, reduction of local chromatin compaction, specific effect on transcriptions of imprinted or X chromosome genes, massive epigenetic changes and alteration of topological organization at the most active chromatin domains (Fan et al. 2005; Maclean et al. 2011; Yang et al. 2013).

H1s in plants are divided into two groups: the main H1s which are ubiquitously and stably expressed, and the minor H1s which are stress-induced and evolutionarily conserved from monocotyledonous to dicotyledonous plants. The drought stress and abscisic acid (ABA)-inducible H1 was initially discovered in wild tomato *Solanum pennellii*, followed by its identification in different organisms, including *Arabidopsis* and *Nicotiana tabacum* (Cohen and Bray 1990; Cohen et al. 1996). Being different from mammalians, *Arabidopsis* H1 is encoded by only three genes: a stress-inducible minor variant *H1.3* (*AT2G18050*) and two main variants *H1.1 (AT1G06760)* and *H1.2 (AT2G30620)* (Ascenzi and Gantt 1997; Gantt and Lenvik 1991; Przewloka et al. 2002). H1.1 and H1.2 are distributed in all vegetative tissues and organs, including leaves, roots, hypocotyls and meristems, and H1.3 is detected in constitutive guard cell-specific tissues and facultative environmentally regulated tissues (Rutowicz et al. 2015).

Plant DNA methylation occurs in CG, CHG and CHH sequence contexts which are primarily catalyzed by DNA methyltransferase 1 (MET1), chromomethylase 3 (CMT3) and the *de novo* methyltransferase (domains rearranged methylases, DRMs) (Law and Jacobsen 2010). The majority of DNA methylation occurs in transposable elements (TEs) in CG, CHG and CHH contexts and is essential to the inhibition of TE activity. Substantial methylation is also found in the bodies of active genes, in which generally occurs in CG context (Law and Jacobsen 2010). In *Arabidopsis*, H1 was known to be involved in DNA methylation and gene transcription. Knockdown of H1 in *Arabidopsis* causes stochastic changes in both hypo- and hyper-DNA methylation in a variety of gene contexts (Wierzbicki and Jerzmanowski 2005). The effect of H1 on DNA methylation has been explained by that DNA METHYLATION 1 (DDM1) plays a role in removing H1 from chromatin to facilitate the access of DNA-methylation machinery (Lyons and Zilberman 2017; Rea et al. 2012; Wierzbicki and Jerzmanowski 2005; Zemach et al. 2013). In addition, H1 and DNA methylation jointly repress TEs and aberrant intragenic transcripts (Choi et al. 2020). The mutation of H1 was known to affect nucleosome density (Choi et al. 2020; Rutowicz et al. 2019; Willcockson et al. 2020). Genome-wide nucleosome maps in yeast, animals and plants revealed that the nucleosome distribution patterns correlate with high-order chromatin organization and transcription levels (Lee et al. 2004; Li et al. 2014; Parnell et al. 2008; Sala et al. 2011; Weiner et al. 2010; Yuan et al. 2005). However, the function of H1 in maintaining high-order chromatin organization is poorly understood in plant.

Hi-C is widely used to map chromatin organization of architectures (Feng et al. 2014; Liu et al. 2016; Wang et al. 2015), and revealed that the chromatin is partitioned into different domains based on different scales and compaction levels, including A/B compartments (Meaburn and Misteli 2007; Wang et al. 2015), TADs (Dixon et al. 2012; Lieberman-Aiden et al. 2009; Nora et al. 2012), and chromatin loops (Jin et al. 2013; Rao et al. 2014). The compartment A for euchromatin with active transcription and compartment B for heterochromatin with repressed transcription can be classified in chromatin regions based on the interaction pattern. There are no obvious TADs, but TAD-like domains in *Arabidopsis* (Dong et al. 2017; Liu et al. 2017). The smaller structure feature is chromatin loops which appear at 10 kb to 1 Mb within TADs (Phillips and Corces 2009) and play roles in transcription, recombination and replication (Mukherjee and Mukherjea 1988). The studies of *Arabidopsis* three-dimensional genome revealed that genome doubling modulates the transcription genome-widely by changed chromatin interactions (Zhang et al. 2019); 3D chromatin organization rearrangement correlates with heat stress-induced transposon activation (Sun et al. 2020); theme of genome structure is the formation of structural units correspond to gene bodies (Liu et al. 2016); KNOT, a structure similar to *flamenco* locus of *Drosophila*, is present in *Arabidopsis* (Grob et al. 2014); and chromatin interactions are related to various epigenetic marks in active or inactive chromatin (Feng et al. 2014).

Recently, partial (about 50 %) depletion of H1 in mouse embryonic stem cells was found to cause no significant change in the overall 3D genome (Geeven et al. 2015); The 3D genome organization is correlated with chromatin compaction and the epigenetic landscape in mouse partial *h1* mutants including *h1c/h1d/h1e* triple (Willcockson et al. 2020) and *h1c/h1e* double mutant (Yusufova et al. 2020). In this study, we generated *Arabidopsis* H1 null mutants, and applied Hi-C and genome-wide approaches to revealing the roles of H1 in regulating 3D genomic organization, nucleosome distribution, DNA methylation and transcription. We showed that H1 depletion impairs the functional links among 3D chromatin interactions, DNA methylation or transcription. We further revealed that H1 modulates the nucleosome density and distribution along the chromosomes, which correlate with the changes of 3D chromatin structure, DNA methylation and transcription.

## Results

### The phenotypes and transcriptome of *Arabidopsis* linker histone *h1* null mutant

To analyze the role of H1 in chromatin organization, we first generated an *h1.1-1h1.2-1h1.3-1* triple T-DNA insertion line which shows no obvious developmental defects (Supplemental Fig. S1A, B) and has only slightly decondensed chromocenters (Supplemental Fig. S1C) as the transcription of *H1.2* in this mutant with T-DNA inserted in the promoter of *H1.2* is significantly induced (Supplemental Fig. S1D). We then constructed *h1* null mutants by crossing *h1.1-1 h1.3-1* double mutant with two CRISPR/Cas9-edited (Feng et al. 2013; Yan et al. 2015) independent *h1.2* mutants, *h1.2-2* and *h1.2-3*, which have pre-mature stop codons due to a nucleotide insertion of “G” or “A” in the second exon of *H1.2*, 949 bp and 906 bp from transcription start site (TSS), respectively (Supplemental Fig. S1E). To eliminate the potential off-target mutations in the CRRISPR experiments, we compared the phenotypes between *h1.1-1 h1.2-2 h1.3-1* (*h1-1*) and *h1.1-1 h1.2-3 h1.3-1* (*h1-2*) with each of them harboring an independent CRISPR/Cas9-edited *h1.2* mutation, we found that these two lines show similar phenotypes including decreased growth and serrated leaves (Fig. 1A, B). For further studies, we used *h1-1* (named as *h1* hereafter) in which the complete deletion of H1 was confirmed by western blots, compared to the partial depletion of H1 in *h1.1-1 h1.2-1 h1.3-1* mutant (Supplemental Fig. S1F). The primary roots of *h1* seedlings are shorter than those of wild type (Col) (Fig. 1*C*), and enlarged nuclei in *h1* leaves were indicated by guard cell nuclei without endoreduplication (Supplemental Fig. S1G, H). At the subnuclear level, we observed that H1 is highly enriched in DAPI-dense heterochromatic chromocenters in wild type (Supplemental Fig. S2A), and depletion of H1 causes dramatic decondensation of chromocenters (Fig. 1D). We then compared the subnuclear distributions of H1.1, H1.2 and H1.3, and found that H1.2 co-localizes with H1.1 or H1.3 in nuclear foci by transiently co-expressing H1.1-YFP/H1.2-mCherry or H1.3-YFP/H1.2-mCherry (Supplemental Fig. S2B).

**Figure 1.**
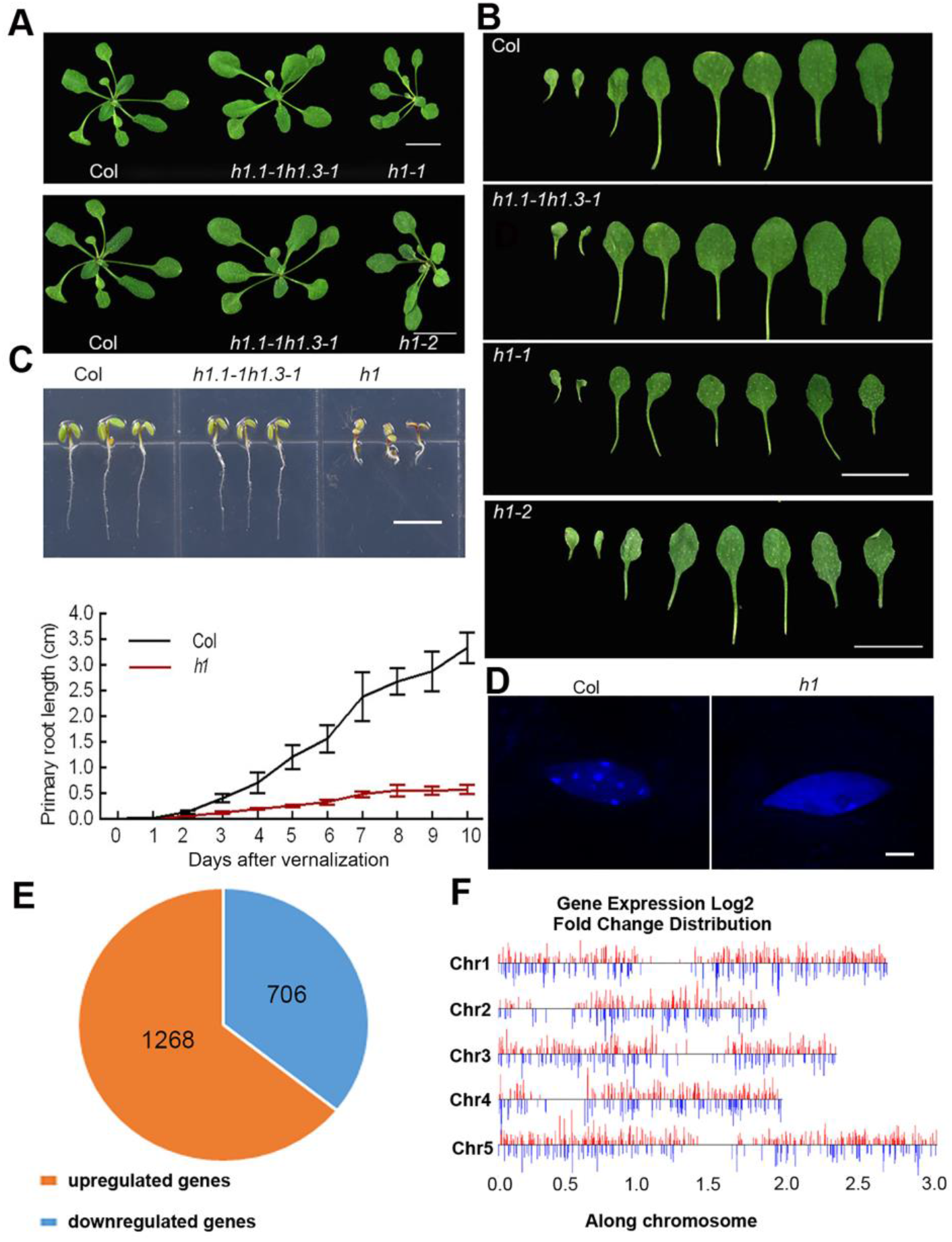
Visual phenotypes and gene expression of *Arabidopsis* seedlings upon H1 depletion. (*A*) Visual phenotypes of *h1-1* and *h1-2* compared to Col and *h1.1h1.3* plants. (Scale bar, 1 cm). (*B*) Rosette leaves of Col, *h1.1h1.3* and *h1* mutant plants. (Scale bar, 1 cm). (*C*) Phenotypes of Col, *h1.1h1.3* and *h1* seedlings grown 4 d after vernalization. (Scale bar, 1 cm), and statistics of primary root lengths of *h1* and Col plants. Error bars represent mean±SDs (n=30). (*D*) Nuclei of *h1* and Col leaf epidermal cells shown by DAPI staining. (Scale bar: 4 μm.). (*E*) Numbers of up-regulated and down-regulated genes (|log2fold change| > 1) in *h1* mutant compared to Col. The data from three biological replicates were combined. (*F*) Distribution of DEGs in genome. The algorithm of DEseq2 and PossionDis were performed to detect the DEGs.

To evaluate the impact of H1 on transcription, we profiled the transcript levels in *h1* by RNA-seq. Among 1974 genes significantly regulated in *h1*, 1268 are up-regulated and 706 down-regulated (Fig. 1E; Supplemental Fig. S3A; Supplemental Table S1). Gene ontology (GO) analysis revealed that these differentially expressed genes (DEGs) are involved in a variety of response processes (Supplemental Fig. S3B; Supplemental Table S2), indicating the functions of H1 in responses to environmental and developmental cues. To verify the results of RNA-seq, we analyzed the transcript levels of several response-related genes by qRT-PCRs. The results were consistent with those in RNA-seq (Supplemental Fig. S3C; Supplemental Table S1). In addition, we found that the differentially up-regulated and down-regulated genes distribute in all five chromosomes (Fig. 1F).

### H1 depletion causes the dampened inter- and intra-chromosomal interactions and extensive A/B compartment switches

To dissect genome-wide chromatin architecture, we performed Hi-C experiments of *h1* mutant (Supplemental Table S3) and Col with each of them having a high reproducibility between two biological replicates (Supplemental Fig. S4A, B). The Hi-C data of Col were obtained from our previous report (Zhang et al. 2019), and used as the control because the seedlings of *h1* null mutant and Col were grown and sampled in parallel at the same time and conditions. Compared to the typical Hi-C heatmap of Col, a clearly homogenous pattern of Hi-C heatmap was observed for *h1* mutant (Fig. 2A). To address the chromosome clustering traits, we calculated the interaction difference matrix from the reads with the same sequencing depth between Col and *h1*. The genome-wide chromatin interaction difference matrix between Col and *h1* revealed that *h1* knockout results in dampened inter- and intra-chromosomal interactions (Fig. 2A-C), decreased inter- or intra-arm interaction (Fig. 2D, E), and inter-pericentromeric or telemetric interaction frequencies (Fig. 2F, G).

**Figure 2.**
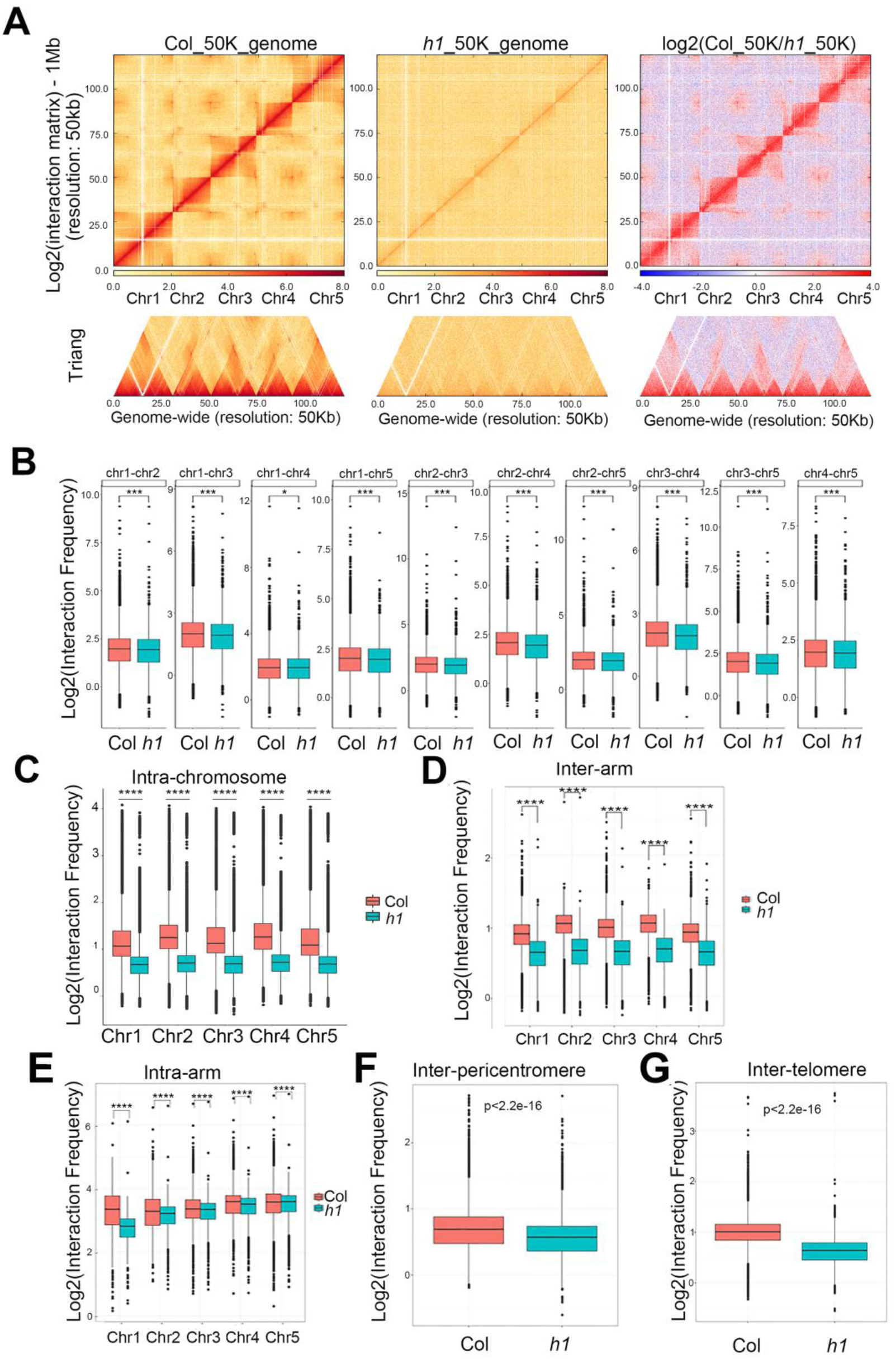
H1 depletion changes chromatin interactions. (*A*) Chromatin interaction heatmaps of Col and *h1*, and differential chromatin interaction heatmap between Col and *h1* at a 50 kb resolution. Chromosomes stacked from bottom left to up right were chr1, chr2, chr3, chr4 and chr5. (*B*) Boxplots showing inter-chromosome interaction frequencies among all chromosome pairs between Col and *h1*. (*C*) Boxplots showing intra-chromosome interaction frequencies between Col (brick red) and *h1* mutant (blue). (*D*) Boxplots showing inter-arm interaction frequencies between Col (brick red) and *h1* mutant (blue). (*E*) Boxplots showing intra-arm interaction frequencies between Col (brick red) and *h1* mutant (blue). (*F*) Boxplots showing pericentromeric interaction frequencies between Col (brick red) and *h1* mutant (blue). (*G*) Boxplots showing telomeric interaction frequencies between Col (brick red) and *h1* mutant (blue). (\*\*\**p*<0.001, \*\**p*<0.01, \**p*<0.05, NS *p*>0.05. The *p* values were tested by Wilcoxon–Mann–Whitney test).

To quantitatively assess the chromatin contacts, we calculated interaction decay exponents (IDEs), which characterize chromatin packing as the slopes of a linear fit of average interaction intensities detected at a given range of genomic distance (Grob et al. 2014). The results displayed that IDEs of intra-chromosome arms (Supplemental Fig. S5A-F), pericentromeres (Supplemental Fig. S6A-F) and telomeres (Supplemental Fig. S7A-F) in *h1* are all lower than those in Col.

Next, we defined the active (A) and inactive (B) chromatin compartments in *h1* and Col by Pearson Correlation (PC1) values (Fransz et al. 2000; Grob et al. 2013). We compared the compartments A/B through the first principal component at a 50 kb resolution between Col and *h1* mutant. We found that 31.0% and 27.5% of the genome showed conserved compartment A and B, respectively (Fig. 3A-C). We observed that H1 depletion induces extensive switches between chromatin compartment A and B (Fig. 3A-C; Supplemental Fig. S8A-D; Supplemental Fig. S9A-D), with 18.5% converted from compartment A to B, and 22.9% converted from compartment B to A (Fig. 3C). In addition, we found that the compartment transition from A to B occurs more than B to A on chromosome 1 (Chr1), Chr2, Chr3 and Chr5, and B to A more than A to B only on Chr4 (Fig. 3D).

**Figure 3.**
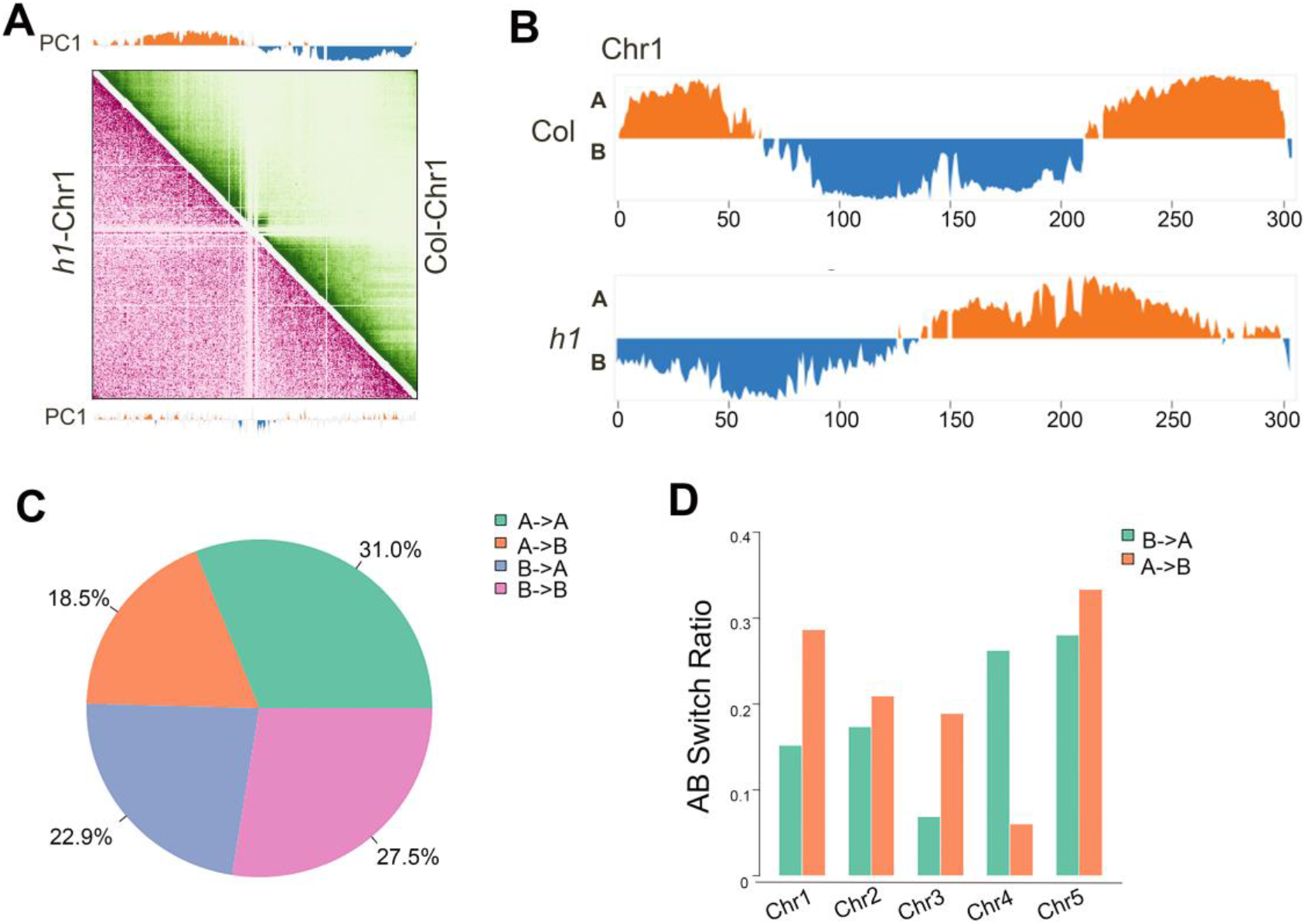
H1 depletion changes chromatin compartmentalization. (*A*) Pearson correlation coefficient matrix and its respective first eigenvector of chr1. The orange regions in the first eigenvector represent compartment A, and the blue regions represent compartment B. Genomic bin size: 50 kb. (*B*) First eigenvector of chr1. Compartment A is presented in orange and compartment B in blue. (*C*) Pie chart representing the percentages of chromatin compartment switches between Col and *h1* mutant. (*D*) Bar graph showing the statistics of structure domain changes in all chromosomes between Col and *h1* mutant.

### The nucleosome density decreases in centromeric regions and increases in chromosome arms in *h1* mutant

To study the potential relationship between the observed weakened chromatin interactions in *h1* and nucleosome density, we first examined micrococcal nuclease (MNase) accessibility which reflects the nucleosome occupancy and gene accessibility by digesting naked DNA through MNase (Li et al. 2014). The results showed that the chromatin in *h1* is more resistant to MNase digestion than Col (Fig. 4A), and all of the mononucleosome ratios were significantly decreased under different MNase levels (Fig. 4B). In addition, western blots indicated that the total H3 and H4 protein levels increase in *h1* compared to Col (Fig. 4C, D). These results indicated that the complete depletion of H1 results in an elevated average density of nucleosomes in chromatin.

**Figure 4.**
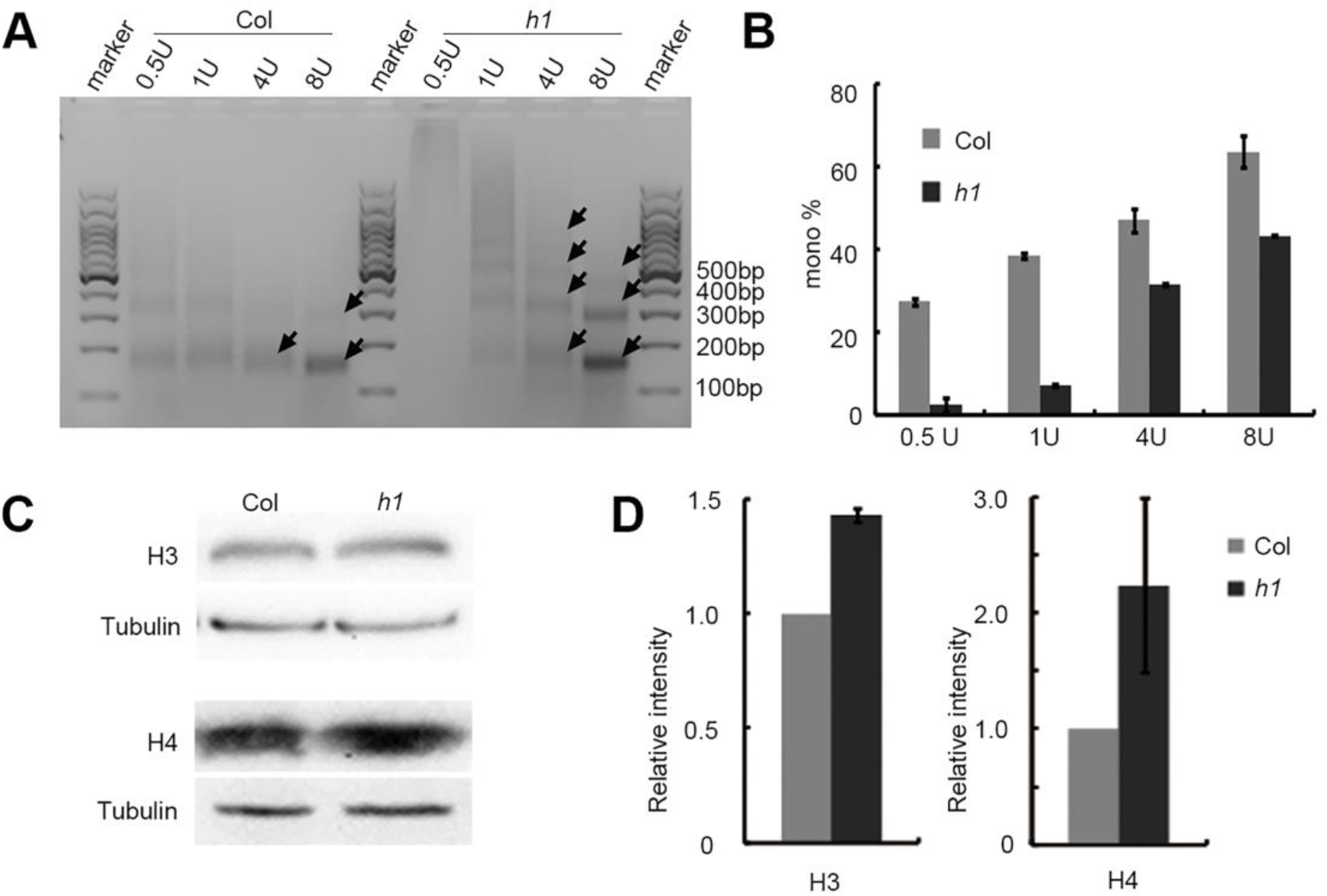
H1 affects the nucleosome density. (*A*) Different titration levels of MNase in the reaction of nuclei extracted from *h1* and Col plants. *h1* is more insensitive to MNase than Col. (*B*) Quantification of mono-nucleosome in *h1* and Col seedlings from the three biological replicates of MNase digestion reaction. The data was analyzed by Gel-Pro analyzer, the mononucleosome ratios are defined as the gray value of the nucleosome monomer band divided by the gray value of the whole lane. (*C*) Western blot on total H3 and H4 in *h1* and Col plants. One of three biological replicates with similar results is shown. (*D*) Quantification of H3 and H4 protein abundance in *h1* and Col seedlings from western blot analysis in three biological replicates.

To examine the chromosomal distribution of nucleosomes in *h1* null mutant, we performed MNase accessibility assays followed by sequencing (MNase-seq) (Supplemental Table S4) in Col and *h1* mutant. The results showed that significant alteration of nucleosome density was presented along all the five chromosomes (Supplemental Fig. S10). To refine the changes of nucleosome packaging in different chromosome regions, we analyzed the nucleosome density on centromeres, pericentromeres and chromosome arms, respectively. We found that the nucleosome density dramatically decreases on centromeres, while increases in both right- and left-chromosome arms. In contrast, nucleosome density in the right- and left-pericentromeres showed no obvious change (Supplemental Fig. S11A-E).

We then analyzed the nucleosome distribution in different types of genes, and found that the overall patterns of nucleosome distribution in protein-coding genes (PCGs), pseudogenes and transposable elements (TEs) in *h1* mutant were similar to those in Col. The evenly spaced nucleosome distribution is observed in gene bodies of PCGs, but not in promoters of PCGs (Supplemental Fig. S12A-C). However, compared to Col, PCGs display a higher nucleosome density in their promoter regions (Supplemental Fig. S12A); pseudogenes and TEs show a slight decrease of nucleosome density (Supplemental Fig. S12B, C).

We further analyzed the nucleosome distribution in up-regulated, down-regulated and unregulated genes in *h1* mutant. We observed that the evenly spaced nucleosome distribution is absent in the bodies of DEGs, but kept in the bodies of non-DEGs upon H1 depletion (Supplemental Fig. S12D-F). In addition, the nucleosome density in *h1* mutant is elevated in promoters, but not in gene bodies, irrelevant of DEGs or non-DEGs (Supplemental Fig. S12G, H).

### The levels of CHG and CHH DNA methylations increase in centromeric regions and decrease in chromosome arms in *h1* mutant

Loss of H1 was reported to cause the elevation of total genomic DNA methylation (Rea et al. 2005). Given that the nucleosome density shows variation among different chromosomal regions, we investigated the potential variations of DNA methylation along chromosomes upon complete depletion of H1. Compared to Col, we found that the DNA methylation level (Supplemental Table S5) in *h1* null mutant increases most obviously over centromeric regions (Supplemental Fig. S13). In addition, DNA methylation levels in CG, CHG and CHH sequence contexts show different distribution patterns along chromosomes (Supplemental Fig. S13B), with the hypermethylation and hypomethylation of CG more evenly distributed in the genome. In contrast, the hypermethylation sites of CHG and CHH locate mainly around centromeres, while the hypomethylation sites of CHG and CHH distribute on chromosome arms with the hypomethylation sites of CHH enriched in the chromosome regions adjacent to centromeres (Supplemental Fig. S13C). Compared to that the nucleosome density decreases in centromeric regions and increases in chromosome arms in *h1* mutant, the reverse correlation between DNA methylation and nucleosome density in centromeric regions and chromosome arms was thus indicated.

Next, we analyzed the effect of H1 on DNA methylation in various DNA elements, and found that the DNA methylation level of repeat sequences increased significantly in *h1* (Supplemental Fig. S14A). In addition, we found that hypermethylation and hypomethylation of gene bodies have higher ratios for longer genes, and hypermethylation of TEs have a higher ratio for longer TEs, while hypomethylation of TEs is not related to their sizes upon *h1* mutation (Supplemental Fig. S14B).

To examine potential genes that might regulate DNA methylation in *h1* mutant, we analyzed the transcription levels of DNA methyltransferases (*MET1*, *DRM2*, *DRM3*, *CMT2* and *CMT3*), DNA demethylase (*DME*, *ROS1*, *DML2* and *DML3*), histone methyltransferases (*SUVH4*, *SUVH5* and *SUVH6*), histone demethylase (*IBM1*) and the chromatin remodeling factor (*DDM1*) involved in DNA methylation directly or indirectly in RNA-seq data. We found that the transcription levels of these DNA methylation-related genes do not change obviously in the absence of histone H1 (Supplemental Fig. S15), implying that the change of DNA methylation might not result from the altered levels of DNA methylation-related genes in *h1* mutant.

### DNA methylation is uncoupled from the transcriptional regulation in *h1* mutant

We then analyzed the relationship between transcription and DNA methylation which generally serves as a repressive marker of transcription (Zemach et al. 2010). To this end, we first analyzed the normalized RNA-seq FPKM of all differential hyper- and hypo-methylated genes, and found that there are no obvious differences (Fig. 5A). We then analyzed the normalized RNA-seq FPKM of DEGs intersected with hyper- or hypo-methylation genes. We found that FPKMs of both hyper- and hypo-DNA methylation genes in *h1* were higher than those in Col (Fig. 5B), in contrast to expectation that the FPKM of hyper-methylation genes decreases and FPKM of hypo-methylation genes increases.

**Figure 5.**
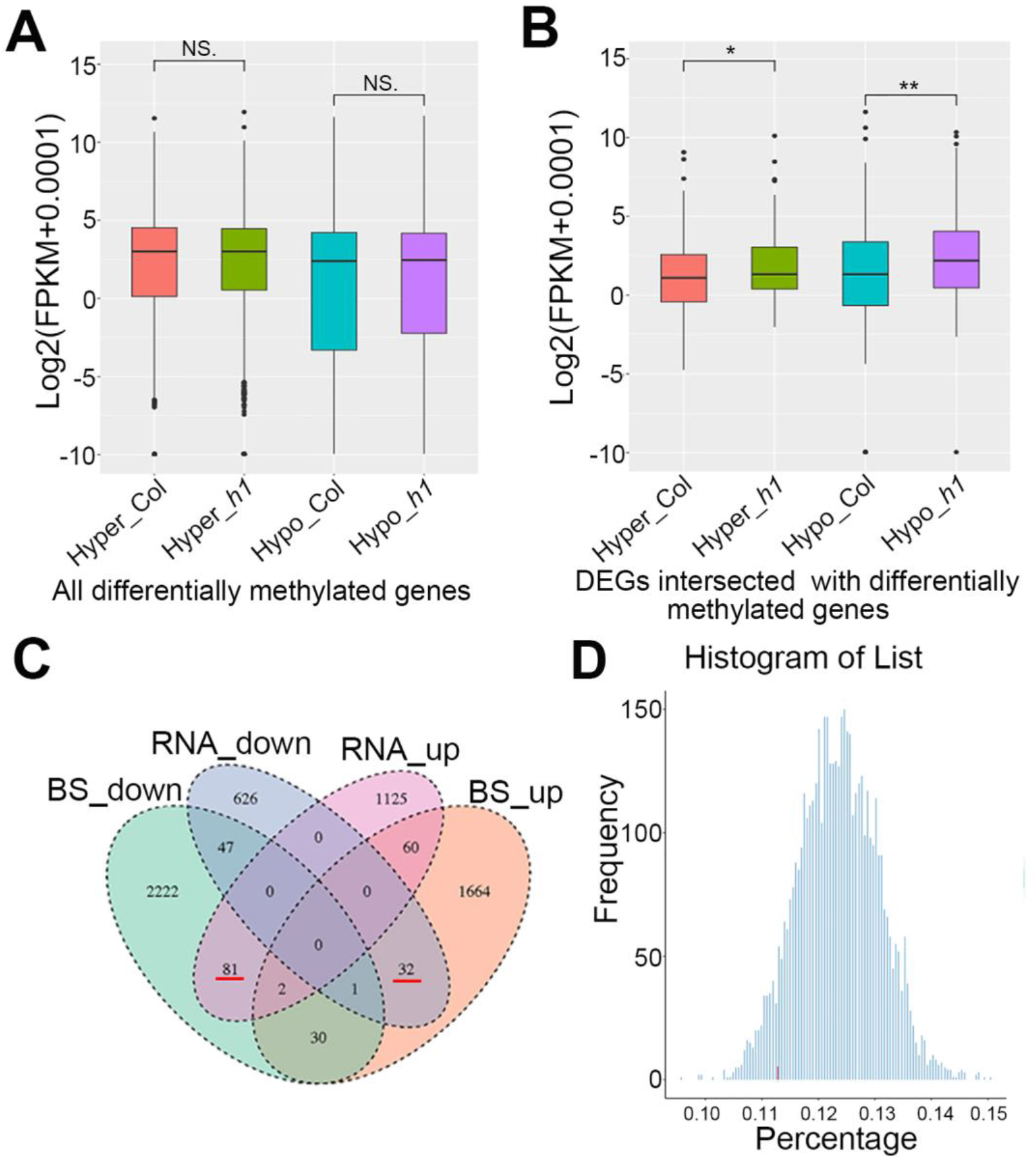
The relationship between DNA methylation and transcription upon H1 depletion. (*A*) Boxplot showing the normalized RNA-seq FPKM of all differentially methylated genes. (*B*) Boxplot showing the normalized RNA-seq FPKM of intersected differentially methylated genes and expressed genes. (\*\*\**p*<0.001, \*\**p*<0.01, \**p*<0.05, NS *p*>0.05. The *p* values were tested by Wilcoxon–Mann–Whitney test). (*C*) Venn diagrams showing numbers of genes WGBS-down (green), RNA-down (blue), RNA-up (pink) and WGBS-up (brick red) in Col and *h1* mutant. (*D*) Histogram of randomly selected no-regulated genes (1974) in differential methylation bins (group number = 5000). The red bar shows the percentage of DEGs in DNA methylation-related genes, and the blue bar chart shows the percentage of equal number (1974) of non-regulated genes in DNA methylation-related genes.

We found that only a small portion of DEGs (233/1974, about 11.29%) localizes in the differential DNA methylation regions (Fig. 5C). Among these 233 genes, 81 genes are RNA-up-regulated and DNA methylation-down-regulated, and 32 genes are RNA-down-regulated and DNA methylation-up-regulated as indicated in Venn diagram which includes RNA-seq data of 1974 DEGs and WGBS data of 4139 differentially methylated genes (Fig. 5C; Supplemental Table S6). Given that DNA methylation can also serve as an active marker in some circumstances (Harris et al. 2018), we then clarified the overall relationship between transcription and DNA methylation by bootstrapping randomized analysis by randomly selecting 5000 group of equal number (1974) of non-regulated genes to determine the percentage of those genes overlapped with DNA methylation-related genes. The result showed that the percentage of DEGs localized in the percentage of randomly selected genes (Fig. 5D), indicating that DNA methylation is no longer a regulatory factor of gene transcription upon H1 depletion.

### DNA methylation is uncoupled from the chromatin compartmentalization in *h1* mutant

DNA methylation was known to correlate with chromatin compartmentalization, and used to reconstruct compartments A/B in Hi-C analysis (Fortin and Hansen 2015). To gain insight into the correlation between DNA methylation and chromatin structure, we analyzed DNA methylation in the chromatin regions with converted compartments. We found that only 20% differentially methylated genes (830/4139, about 20%) overlapped with chromatin compartment A/B conversions (Fig. 6A). In addition, we found the Cs, CG, CHG, and CHH methylation levels of both A to B and B to A chromatin compartment transition-related genes in *h1* are lower than those in Col (Supplemental Fig. S16A-D), in contrast to the general expectation that the methylation level in A to B regions increases, while that in B to A regions decreases.

**Figure 6.**
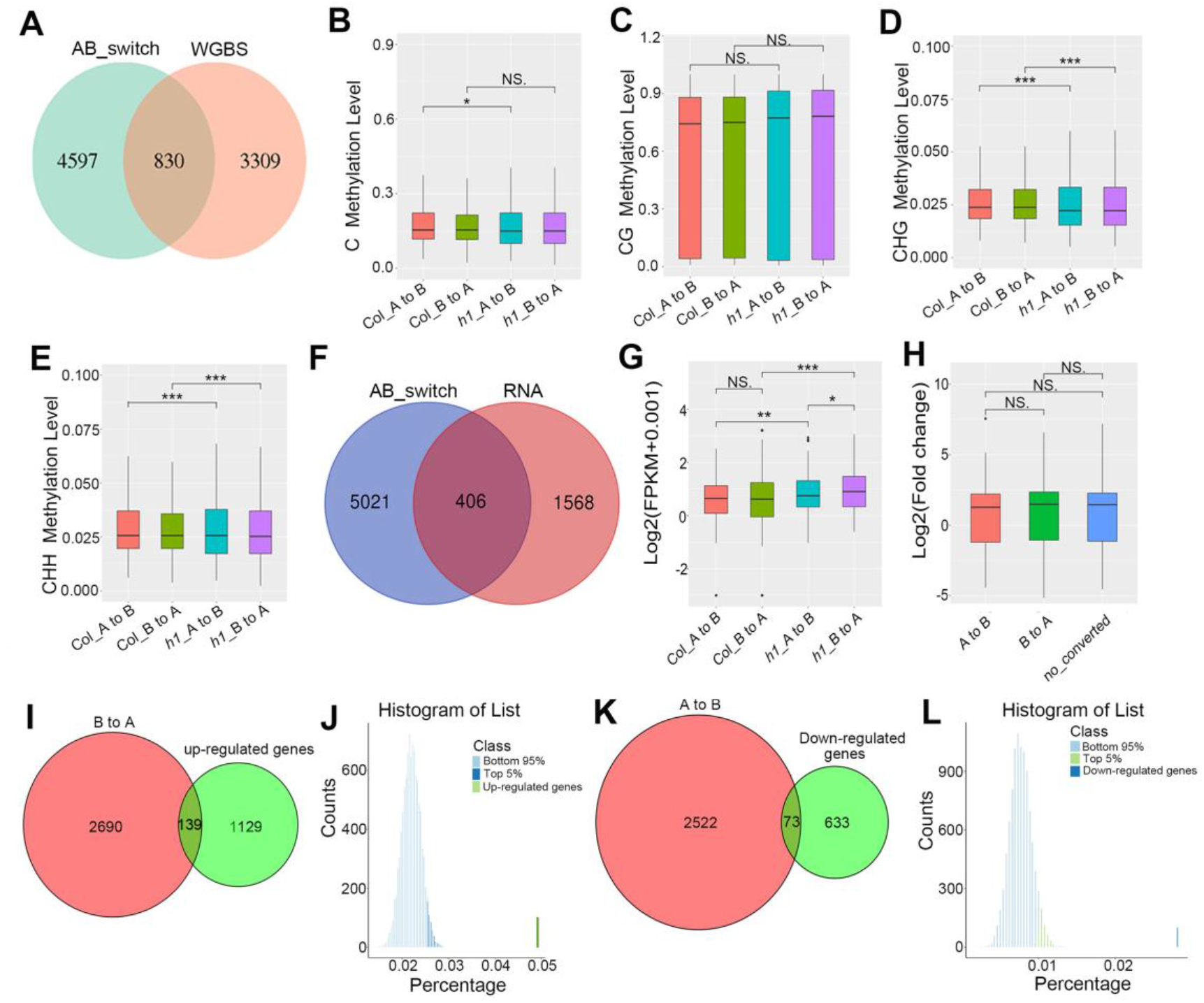
The relationships between chromatin compartments A/B switches and transcription or DNA methylation in *h1* mutant. (*A*) Venn diagram showing number of A/B switched genes (blue) and WGBS differential genes (pink) in *h1* compared to Col. (*B*-*E*) Boxplot showing all Cs (*B*), CG (*C*), CHG (*D*), or CHH (*E*) types of methylation levels in intersected genes between compartment A/B switches and DNA methylation, respectively. (*F*) Venn diagrams showing numbers of genes in switched regions between compartment A/B (dark blue) and DEGs (pink) in Col and *h1* mutant. (*G*) Boxplot showing the transcription levels in Col and *h1* in the switched chromatin regions between compartment A and B. (*H*) Boxplot showing log2 fold change of expression levels of genes in regions with switched chromatin domain between compartment A and B or without compartment transition (no converted). (*I*) Venn diagrams showing numbers of genes in compartment B to A (pink) and up-regulated genes (green) in *h1* mutant compared to Col. (*J*) Histogram of randomly selected no-regulated genes (139) in differential interaction bins (group number = 5000). The top 5 percentile of randomly selected control genes was labeled in blue. (*K*) Venn diagrams showing numbers of genes in compartment A to B (pink) and down-regulated genes (green) in *h1* mutant compared to Col. (*L*) Histogram of randomly selected down-regulated genes (73) in differential interaction bins (group number = 5000). The top 5 percentile of randomly selected control genes was labeled in green. (\*\*\**p*<0.001, \*\**p*<0.01, \**p*<0.05, NS *p*>0.05. The *p* values were tested by Wilcoxon–Mann–Whitney test).

We then analyzed the DNA methylation levels of genes intersected between DNA methylation and A/B switches, and found that the Cs, CG, CHG, and CHH methylation levels of both compartments A to B or B to A switch-related genes in *h1* are lower than those in Col (Fig. 6B-E). We further analyzed the promoter regions and TSSs of genes intersected between DNA methylation and compartment A/B switches (Supplemental Table S7). The results indicated that the Cs, CG, CHG, and CHH methylation levels of both compartments A to B or B to A switch-related genes in *h1* are lower than those in Col (Supplemental Fig. S16E-L). Together, we concluded that there is no correlation between DNA methylation and chromatin compartment A/B conversions in *h1* mutant.

### The switches of compartment A/B are largely uncorrelated with gene transcription

To reveal the role of chromatin folding in regulating gene expression upon H1 depletion, we analyzed the correlation between chromatin compartment switches and transcription. We found that most of DEGs (1568/1974, about 79.4%) show no overlap with chromatin compartment switches between A and B (Fig. 6F) with only a small portion of DEGs (406/1974, about 20.6%) overlapped with chromatin compartment A/B switches (Fig. 6F). In addition, although chromatin compartment switches account for 41.4% (18.5%+22.9%, Fig. 3C) of the *Arabidopsis* genome, only 7.7% (1974/25498) (Arabidopsis Genome Initiative 2000) of genes are DEGs in *h1* mutant, supporting that most of the A/B switches do not affect transcription.

To further evaluate the relevance between A/B switches and transcription, we analyzed the normalized RNA-seq FPKM of genes in compartment A/B switches, and found that genes in compartments transited from B to A show only slightly higher expression levels than those in compartments transited from A to B in *h1* mutant (Fig. 6G). We also noticed that most of DEGs (1568 genes) show no overlap with A/B switches (Fig. 6H). We further found that only 10.9 % (139/1268) up-regulated genes localized in compartment B to A switches (Fig. 6I), and 10.3% (73/706) down-regulated genes localized in compartments A to B switches (Fig. 6K). To know the confidence of these results, we performed bootstrapping randomized analysis. We randomly selected 5000 group of equal number (139) of up-regulated genes to determine the percentage of those genes overlapped with compartment A/B switches-related genes. The results showed that the top 5 percentile of randomly selected genes (2.6%) are only slightly lower than the percentage of up-regulated genes localized in compartment B to A switches (139/2690≈5.2%) (Fig. 6J). In addition, we randomly selected 5000 group of equal number (73) of down-regulated genes to determine the percentage of those genes overlapped with compartment A/B switches-related genes, and found that the top 5 percentile of randomly selected genes (1.2%) are only slightly lower than the percentage of up-regulated genes localized in compartments A to B switches (73/25220 ≈2.9%) (Fig. 6L). We concluded that chromatin compartment switches are predominantly not related to the gene regulation in *h1* knockout mutant.

## Discussion

The linker histone H1 binds and locks the adjacent nucleosomes to maintain chromatin structure (Fan et al. 2005). In the study, we found that unlocking in and out ends of nucleosome DNA by H1 complete depletion dampens the chromatin interactions, alters nucleosome density and distribution, changes the level and distribution of DNA methylation and affects the expressions of a subset of genes. Notably, loss of H1 disrupts the connections among chromatin interactions, DNA methylation and transcription, in which the H1 depletion-induced changes of nucleosome density and distribution might play an important role.

### The role of linker histone H1 in regulating plant growth and development

*Arabidopsis*, which harbors only three genes encoding linker histone H1, is a good model organism to study the functions of H1. The partial mutants were reported, such as RNAi triple knockdown mutant (Wierzbicki and Jerzmanowski 2005), or T-DNA insertion triple knockdown mutant *h1.1-1 h1.2-1 h1.3-1* with subtle phenotypes at developmental transitions (Rea et al. 2012; Rutowicz et al. 2019). In this study, we generated knockout mutants of histone *H1s* by T-DNA insertions and CRISPR/Cas9-based gene editing, which show pleiotropic phenotypes and are fertile. RNAi lines of H1 showed pleiotropic developmental abnormalities including changed size, serrated, small or elongated leaves, and reduced apical dominance, and plants with aberrant phenotypes have a considerably greater reduction of H1 expression than plants with no visible changes (Wierzbicki and Jerzmanowski 2005). We observed the *h1* null mutant phenotypes of small plant size, short primary roots, serrated, small and elongated leaves, indicating the roles of H1 in plant growth and development by regulating a subset of genes.

There are 11 H1-coding genes in human and mouse, including replication dependent *H1.1-H1.5*, non-replication dependent *H1.0* and *H1.X*, sperm specific *H1t*, *H1T2* and *H1LS1*, and egg specific *H1oo* (Godde and Ura 2009; Kowalski and Palyga 2012). Knockouts of *H1.3*, *H1.4* and *H1.5* in mouse resulted in the death of embryos with altered distribution of nucleosomes and without change of the nuclear size (Fan et al. 2003). However, *Tetrahymena* strains that lacked either macronuclear or micronuclear histone H1 proteins or both were fully viable, showing similar growth to the wild type strains, implicating that H1 is more important for higher organisms. The increasing number of *H1* genes during the evolution of higher organisms might be due to the needs for precise and specific regulation of gene expressions or responses to environmental stimuli as the tissue-specific expression of H1 was observed in mammalian and the expressions of response genes were induced by H1 depletion as shown in this study. H1 can also affect the phenotypes of organisms by modulating the post-translational modifications of histones (Ausio 1992; Herrera et al. 2000).

### The role of H1 in functional connections among chromatin interactions, DNA methylation and transcription

At the cellular level, H1 depletion results in the increased nuclear size and decondensation of heterochromatic chromocenters. Hi-C analysis showed that the complete depletion of H1 in *Arabidopsis* dampens the chromatin interactions, supporting the role of linker H1 in the high-order chromatin structure in addition to its role in facilitating the folding of chromatin into 30 nm fiber. Moreover, significant alterations of nucleosome density and distribution were induced along all chromosomes. The nucleosome repeat length of H1-rich genes was observed to decrease substantially in *Arabidopsis h1* knockdown mutants (Choi et al. 2020). The increase of nucleosome density and decrease of nucleosome repeat length might prevent chromatin DNA from getting closer to each other, leading to increased nuclear size and decreased chromatin interactions which might affect the nucleating of heterochromatin in chromocenters.

Further, we found the extensive switches between chromatin compartment A and B upon H1 depletion, indicating the important function of H1 in defining the chromatin domain identity. A/B compartments were known to be related to DNA methylation (Fortin and Hansen 2015). However, our results showed that both the changes of DNA methylation and expressions of most DEGs are uncorrelated with the chromatin compartment switches in *h1* mutant, indicating that the linker histone also serve as a linker between chromatin compartment identity and DNA methylation directly or indirectly. Our results from the *Arabidopsis* H1 knockout line are different from those in mammalian or human H1 knockdown cells. In embryonic stem cells with about 50 % depletion of H1, the 3D genome showed no significant change and the alterations in TAD configuration coincide with epigenetic landscape changes but not with transcriptional output changes (Geeven et al. 2015). In addition, most A/B genomic compartments and TADs are unchanged upon deletion of H1C, H1D and H1E in mammals (Willcockson et al. 2020). In *h1c/h1d/h1e* triple and *h1c/h1e* double mutants, the chromatin architectural and epigenetic changes underlie the transcriptional alterations (Willcockson et al. 2020; Yusufova et al. 2020). How H1 maintains the relationship between the 3D structure of chromatin and DNA methylation is worthy of further study.

Interestingly, the compartment switches from A to B are more than B to A on chromosome 1 (Chr1), Chr2, Chr3 and Chr5, while B to A are more than A to B only on Chr4 (Fig. 3D), which might be related to the specific chromatin status of Chr4 with nucleolar associated chromatin domain (NAD) and nucleolar organizer region 4 (NOR4) (Pontvianne et al. 2016; Rabanal et al. 2017).

### The correlation between the nucleosome density and DNA methylation

In eukaryotes, DNA methylation plays important roles in chromatin structure (Zhang et al. 2017). In ascomycete fungi, loss of H1 leads to genome-wide hypermethylation (Barra et al. 2000; Seymour et al. 2016). In mouse, the mutation of H1 reduces the DNA methylation levels in specific loci (Yang et al. 2013). In *Arabidopsis*, knockdown of H1 increases DNA methylation levels of heterochromatic elements and decreases DNA methylation levels of euchromatic TEs in all sequence contexts (Choi et al. 2020; Rutowicz et al. 2015; Zemach et al. 2013). In animals, CpG methylation induces tight wrapping of DNA around the histone core accompanied by a topologic change, and the changes in physical properties of nucleosomes induced by CpG methylation may contribute to the formation of repressive chromatin architectures (Lee and Lee 2012). The non-CG methylations (CHG and CHH) in plant are mainly distributed in heterochromatin region (Du et al. 2015). Our results revealed that the levels of CHG and CHH DNA methylations increase in centromeric regions and decrease in chromosome arms, while the nucleosome density dramatically decreases in centromeres and increases in chromosome arms in *h1* mutant. One possible reason might be that the more nucleosome facilities stronger chromatin condensation, which prevents DNA methyltransferases or DNA demethylases to access to target sites as the transcription levels of these DNA methyltransferase or demethylase genes do not change obviously. The study of DDM1 supported this idea, showing DDM1 remove H1 from chromatin in order to facilitate the access of DNA methylation factors (Lyons and Zilberman 2017). Other studies also pointed out that nucleosomes are substantial obstacles to DNA methylation (Baubec et al. 2015; Huff and Zilberman 2014). However, the precise mechanism of H1 involved in DNA methylation needs to be intensely investigated, and the functions of HMGA proteins, which compete with histone H1 to bind to linker DNA (Catez and Hock 2010; Charbonnel et al. 2014), also need to be examined upon H1 complete depletion.

### The role of H1-defined nucleosome distribution in the gene expression

Only a subset of genes are specifically regulated upon depletion of H1, however, the underlying mechanism about the specific gene regulation through H1 is not clear. Upon unlocking in and out ends of nucleosome DNA by H1 complete depletion, the nucleosome density in promoters elevates, but not in gene bodies (Supplemental Fig. S12G, H), and the phased distribution pattern of nucleosome in gene bodies of regulated genes is lost, but not in gene bodies of unregulated genes (Fig. 5D, F), indicating the role of the specific nucleosome deposition defined by H1 in the regulation of gene expression. Alternatively, transcription may also affect the differential nucleosome distribution between promoter and gene bodies. It was reported that H1 inhibits RNA polymerases from binding to chromatin (Krishnakumar et al. 1995), and regulates neuronal activation through modulating immediate early gene (IEG) expression, in which H1 is replaced by PARP on IEG promoters after neuronal stimulation (Azad et al. 2018). Therefore, H1 may also modulate the specific nucleosome density to regulate accessibility of transcription machinery to chromatin and affect gene expression.

This study unveiled for the first time the effect of H1 on the relationships among the 3D chromatin structure, DNA methylation and gene expression. In animal cells, no obvious change in the three-dimensional chromatin structure was observed upon partial depletion of linker histones (Geeven et al. 2015; Willcockson et al. 2020; Yusufova et al. 2020). In *Arabidopsis*, we observed enlarged nuclei and obvious decondensation of heterochromatic chromocenters, which correlates with a more homogenous pattern of chromatin interaction heatmap and dramatic changes of chromatin compartments upon H1 complete depletion. Our results shed light on the roles of linker histones in the maintenance of proper genome folding and coordinating chromatin compartmentalization, DNA methylation and gene expression.

## Methods

### Plant materials and growth conditions

Wild type *Arabidopsis thaliana* (Col-0 ecotype) and mutant plants were grown under 16 h light/8 h dark at 22 ℃. T-DNA insertion lines of *h1.1-1* (SALK_128430C), *h1.2-1* (SALK_002142) and *h1.3-1* (SALK_025209) were obtained from *Arabidopsis* Biological Resource Center and confirmed by PCR-based genotyping. Primers used were listed in Supplemental Table S8.

The knockout mutants of *h1-1* and *h1-2* through CRISPR/Cas9 -mediated editing were generated as previously reported (Feng et al. 2013; Yan et al. 2015). The sgRNAs were listed in Supplemental Table S8.

### Constructs and transient expression

The cDNAs of *H1.1*, *H1.2*, and H*1.3* were amplified by PCR from Col cDNAs using primers listed in Supplemental Table S8, digested by *Eco*RI/*Sal*I, and subcloned into *Eco*RI/*Sal*I-treated vectors of pCambia1300-35S-N1-YFP and pCambia1300-35S-N1-mCherry (Fang and Spector 2007). The constructs were confirmed by sequencing and introduced into *Agrobacterium tumefaciens* (GV3101) by electroporation.

*Arabidopsis* plants were transformed by the floral dip method (Clough and Bent 1998). Transient expression and colocalization analysis were performed as described (Fang and Spector 2010).

### Western blot

Total proteins were extracted from 10-day-old seedlings. 100 mg materials were collected and ground in liquid nitrogen. The powders were suspended in 3×SDS buffer (sample: buffer 1:3), then boiled at 100 ℃ for 10 min and centrifuged for 2 min at 12,000 rpm, the supernatants were separated on 15 % SDS-PAGE. Antibodies to H3 (Sigma H9289; 1:3000), H4 (Active Motif 61300; 1:3000), and H1 (Agrisera AS111801; 1:3000) were used.

### DAPI staining

Leaves from 4-week-old plants were used. After washes in PBS, nuclei were counterstained in Fluoromount mounting media with DAPI (4′, 6-diamidino-2-phenylindole, YEASEN 36308ES11). Images of nuclei were acquired as described (Shi et al. 2011).

### Hi-C library preparation

Hi-C experiments were performed essentially as described (Grob et al. 2014) with some modifications. Two biological replicates of Col we reported previously (Zhang et al. 2019) (GSE114950) and two biological replicates of *h1* null mutant were sampled in parallel with Col under same growth condition. Briefly, 2.5 g aerial parts of the seedlings were fixed and ground into powder in liquid nitrogen. 600 U *Hin*dIII restriction enzyme were used to digest the extracted nuclei by incubating overnight at 37℃, then the digested chromatin was blunt-ended with 1μl 10mM dATP, dTTP, dGTP and 25μl 0.4mM biotin-14-dCTP and 100 U Klenow fragment for 45 min at 37℃. The ligation reaction was performed in 10 time volume of ligation buffer under constant shaking with 745μl 10× ligation buffer, 10% Triton X-100, 80μl 10 mg/ml BSA and ATP, 100 Weiss U T4 DNA ligase, at 16℃ for 6 h. The nuclei were then reverse-crosslinked with proteinase K at 65℃overnight. Subsequently, the extracted chromatin was fragmented into a mean size of 300 bp using a sonicator (Covaris S220). Hi-C libraries were constructed with NEB Next Multiplex Oligos kit and KAPA Hyper Prep Kit. The final library was subject to sequencing on an Illumina HiSeq 2000 instrument with 2 × 150-bp reads.

### Hi-C sequencing data processing

Hic-pro (Servant et al. 2015) and Bowtie2 (Langmead and Salzberg 2012) were used for Hi-C read mapping. The clean Hi-C reads of Col and *h1* mutant were aligned to *Arabidopsis* reference genome (TAIR10) after removing the adapter. Following with HiC-Pro and Juicer software (Durand et al. 2016), valid pairs of Col and *h1* mutant were used to create interaction matrixes with bin size 50 kb for further analysis. The interaction matrixes were normalized with KR method from Juicer (Durand et al. 2016). The reproducibility of two biological replicates was tested with Pearson correlation coefficient from the interaction matrixes (Lin et al. 2018). After excluding the pericentromeres as reported (Grob et al. 2014), the first principal component was used to identify compartments A and B with Juicer.

### Calculation of chromatin interaction and interaction decay exponents

The normalized interaction matrix from Col was divided by the normalized interaction matrix from *h1* mutant, with all the zeros in matrixes replaced with 1% quintiles from the non-zero elements in each matrix, which were used to analyze the difference of interaction matrixes between Col and *h1* mutant. We used Log2 transformation and median normalization to standardize the difference matrix. Interaction decay exponents (IDEs) were calculated (Grob et al. 2014) for chromosomes, pericentromeres and telomeres to study the interaction frequency changes dependent on the genome distance.

### Bootstrapping analysis

In the bootstrapping strategies (Zhang et al. 2019; Buonaccorsi et al. 2018), we randomly selected 5000 groups (n = 5000 times) of the relevant genes of equal number, which were subjected to the same analysis to determine the percentage of those groups fallen in differential interaction bins or differential methylation bins. The percentile of the test sample lie above the top 5 percentile of the control distribution was considered confidently.

### Micrococcal nuclease treatment assay

MNase assay was performed using protocol described (Zhang et al. 2018) with slight modification. For each MNase assay, 2 g of 10-day-old seedlings were used for nuclear extraction. The materials were ground in liquid nitrogen and shaked in lysis buffer containing 50 mM HEPES (pH 7.5), 10 % glycerol, 1 % Triton X-100, 1mM EDTA (pH 7.5), 150 mM NaCl, 5 mM β-mercaptoethanol and protease inhibitor cocktail (Roche) for 1 h at 4℃. The lysis mixture was filtered through a 40 μm cell strainer (BD) into fresh 50 ml tubes, nuclei were collected by centrifugation at 4℃ for 20 min at 4000 g. Nuclei were then washed twice with buffer A containing 25 mM Tris–HCl (pH 7.5), 0.44 M sucrose, 10 mM MgCl_2_, 0.1% Triton-X and 10 mM β-mercaptoethanol, and washed once with buffer B containing 20 mM Tris–HCl (pH 7.5), 352 mM sucrose, 8 mM MgCl_2_, 0.08% Triton-X, 8 mM β-mercaptoethanol and 20% glycerol, then resuspended in 100 μl of buffer B and flash frozen in liquid nitrogen. The nuclei obtained were divided into four portions for MNase treatments by incubating with 0.5 U, 1 U, 4 U and 8 U of MNase (TaKaRa 2910A) in MNase digestion buffer containing 20 mM Tris–HCl (pH 8.0), 5 mM NaCl and 2.5 mM CaCl_2_ for 20 min at 37℃. Reactions were stopped by addition of 500 mM EDTA. Digested DNA was purified and separated on 2 % agarose gel. For MNase-seq, all the nuclei were treatment with 4 U of MNase and mononucleosome-sized fragments were gel purified for sequencing library generation.

### MNase-seq analysis

Approximately 1 μg of purified mononucleosome-sized DNA fragment was used for Illumina library generation per manufacturer’s instruction. Library construction and sequencing were performed by Genergy Biotechnology Co. Ltd. (Shanghai, China). Sequencing was carried out as single-end 50 bp reads on Illumina HiSeq-2000. Data analysis was carried out as previously described (Li et al. 2014). Briefly, the quality of raw reads was examined by FastQC (http://www.bioinformatics.babraham.ac.uk/projects/fastqc/). All clean reads were mapped to the TAIR 10 genome with the BOWTIE (http://bowtie.cbcb.umd.edu). Nucleosome positions were identified using the TEMPLATE FILTER software (http://compbio.cs.huji.ac.il/NucPosition/TemplateFiltering/Home.html).

### RNA-seq analysis

Total RNAs were extracted from 10-day-old seedlings using the RNeasy plant mini kit (Qiagen). Yield and RNA integrity were detected by using an Agilent 2100 Bioanalyzer, and RNA purity was determined by using a Nanodrop ND-1000 spectrophotometer. cDNA library construction and sequencing were performed by Beijing Genomics Institute (BGI) using BGISEQ-500 platform for 50 bp single-end sequencing as previously described (Huang et al. 2018). At least 20 M clean reads of sequencing depth were obtained for each sample. Three independent biological replicates were performed. The raw reads were trimmed and quality controlled by Trimmomatic with default parameters (http://www.usadellab.org/cms/uploads/supplementary/Trimmomatic). Then clean reads were separately aligned to *Arabidopsis thaliana* genome from TAIR 10 with orientation mode using tophat software (http://tophat.cbcb.umd.edu/). The level for each transcript was calculated using the fragments per kilobase of exon per million mapped reads (FPKM) method. Cuffdiff (http://cufflinks.cbcb.umd.edu/) was used for differential expression analysis. GO functional enrichment analysis was carried out by Goatools (https://github.com/tanghaibao/Goatools). The differential expression analysis was run using the classical normalization method DESeq2 R package (Love Huber and Anders 2014) with a 0.05 *p*-value, 0.05 false discovery rate, and cutoff of 1 log-fold change. The hypergeometric test was performed as previously described (Wollmann et al. 2017).

### DNA methylation analysis

For whole genome bisulfite sequencing (WGBS), genomic DNA (gDNA) was extracted from 10-day-old seedlings with the DNeasy plant mini kit (Qiagen) per manufacturer’s introduction. Library construction and sequencing were performed by Beijing Genomics Institute (BGI) using Illumina HiSeq-2000 for 100 bp paired-end sequencing. The raw paired end reads were trimmed and quality controlled by SeqPrep (https://github.com/jstjohn/SeqPrep) and Sickle (https://github.com/najoshi/sickle) with default parameters. All clean reads were mapped to the TAIR 10 genome with the BSMAP aligner (http://code.google.com/p/bsmap/) allowing up to 2 mismatches. Uniquely mapped reads were used to determine the cytosine methylation levels as previously stated (Lister, O’Malley, Tonti-Filippini, Gregory, Berry, Millar and Ecker 2008).

### Data access

The Hi-C, WGBS and RNA-seq datasets have been submitted to NCBI (PRJNA680865), and MNase-seq datasets have been submitted to NCBI (PRJNA695028).

## Competing interest statement

The authors declare no competing interests.

## Acknowledgments

This work was supported by National Natural Science Foundation of China (31871230 to Y.F. and 31971334 to Z.S.).

## Author contributions

Z.S. and Y.F. designed and performed experiments, analyzed data, and wrote the manuscript. M.L. generated *h1* mutants, acquired DNA methylation and RNA-seq data. Y.W. helped data analysis. H. Z. performed Hi-C experiments.

## Notes

### Competing Interest Statement

The authors have declared no competing interest.

## References

Arabidopsis Genome Initiative. 2000. Analysis of the genome sequence of the flowering plant *Arabidopsis thaliana*. Nature 408: 796–815.

Ascenzi, R. and J.S. Gantt. 1997. A drought-stress-inducible histone gene in *Arabidopsis thaliana* is a member of a distinct class of plant linker histone variants. Plant Mol Biol 34: 629–641.

Ausio, J. 1992. Structure and dynamics of transcriptionally active chromatin. J Cell Sci 102 (Pt 1): 1–5.

Azad, G.K., K. Ito, B.S. Sailaja, A. Biran, M. Nissim-Rafinia, Y. Yamada, D.T. Brown, T. Takizawa, and E. Meshorer. 2018. PARP1-dependent eviction of the linker histone H1 mediates immediate early gene expression during neuronal activation. J Cell Biol 217: 473–481.

Barra, J.L., L. Rhounim, J.L. Rossignol, and G. Faugeron. 2000. Histone H1 is dispensable for methylation-associated gene silencing in Ascobolus immersus and essential for long life span. Mol Cell Biol 20: 61–69.

Baubec, T., D.F. Colombo, C. Wirbelauer, J. Schmidt, L. Burger, A.R. Krebs, A. Akalin, and D. Schubeler. 2015. Genomic profiling of DNA methyltransferases reveals a role for DNMT3B in genic methylation. Nature 520: 243–247.

Bednar, J., R.A. Horowitz, S.A. Grigoryev, L.M. Carruthers, J.C. Hansen, A.J. Koster, and C.L. Woodcock. 1998. Nucleosomes, linker DNA, and linker histone form a unique structural motif that directs the higher-order folding and compaction of chromatin. Proc Natl Acad Sci U S A 95: 14173–14178.

Buonaccorsi, J.P. G. Romeo and M. Thoresen. 2018. Model-based bootstrapping when correcting for measurement error with application to logistic regression. Biometrics 74: 135–144.

Catez, F. and R. Hock. 2010. Binding and interplay of HMG proteins on chromatin: lessons from live cell imaging. Biochim Biophys Acta 1799: 15–27.

Charbonnel, C., O. Rymarenko, I.O. Da, F. Benyahya, C.I. White, F. Butter, and S. Amiard. 2018. The Linker Histone GH1-HMGA1 Is Involved in Telomere Stability and DNA Damage Repair. Plant Physiol 177: 311–327.

Choi, J., D.B. Lyons, M.Y. Kim, J.D. Moore, and D. Zilberman. 2020. DNA Methylation and Histone H1 Jointly Repress Transposable Elements and Aberrant Intragenic Transcripts. Mol Cell 77: 310–323.

Clough, S.J. and A.F. Bent. 1998. Floral dip: a simplified method for Agrobacterium-mediated transformation of *Arabidopsis thaliana*. Plant J 16: 735–743.

Cohen, A. and E.A. Bray. 1990. Characterization of three mRNAs that accumulate in wilted tomato leaves in response to elevated levels of endogenous abscisic acid. Planta 182: 27–33.

Cohen, A., A.L. Plant, M.S. Moses, and E.A. Bray. 1991. Organ-Specific and Environmentally Regulated Expression of Two Abscisic Acid-Induced Genes of Tomato: Nucleotide Sequence and Analysis of the Corresponding cDNAs. Plant Physiol 97: 1367–1374.

Dixon, J.R., S. Selvaraj, F. Yue, A. Kim, Y. Li, Y. Shen, M. Hu, J.S. Liu, and B. Ren. 2012. Topological domains in mammalian genomes identified by analysis of chromatin interactions. Nature 485: 376–380.

Dong, P., X. Tu, P.Y. Chu, P. Lu, N. Zhu, D. Grierson, Du B, P. Li, and S. Zhong. 2017. 3D Chromatin Architecture of Large Plant Genomes Determined by Local A/B Compartments. Mol Plant 10: 1497–1509.

Drabent, B., P. Saftig, C. Bode, and D. Doenecke. 2000. Spermatogenesis proceeds normally in mice without linker histone H1t. Histochem Cell Biol 113: 433–442.

Du J, L.M. Johnson, S.E. Jacobsen, and D.J. Patel. 2015. DNA methylation pathways and their crosstalk with histone methylation. Nat Rev Mol Cell Biol 16: 519–532.

Durand, N.C., M.S. Shamim, I. Machol, S.S. Rao, M.H. Huntley, E.S. Lander, and E.L. Aiden. 2016. Juicer Provides a One-Click System for Analyzing Loop-Resolution Hi-C Experiments. Cell Syst 3: 95–98.

Fan, Y., A. Sirotkin, R.G. Russell, J. Ayala, and A.I. Skoultchi. 2001. Individual somatic H1 subtypes are dispensable for mouse development even in mice lacking the H1(0) replacement subtype. Mol Cell Biol 21: 7933–7943.

Fan, Y., T. Nikitina, E.M. Morin-Kensicki, J. Zhao, T.R. Magnuson, C.L. Woodcock, and A.I. Skoultchi. 2003. H1 linker histones are essential for mouse development and affect nucleosome spacing in vivo. Mol Cell Biol 23: 4559–4572.

Fan, Y., T. Nikitina, J. Zhao, T.J. Fleury, R. Bhattacharyya, E.E. Bouhassira, A. Stein, C.L. Woodcock, and A.I. Skoultchi. 2005. Histone H1 depletion in mammals alters global chromatin structure but causes specific changes in gene regulation. Cell 123: 1199–1212.

Fang, Y. and D.L. Spector. 2007. Identification of nuclear dicing bodies containing proteins for microRNA biogenesis in living *Arabidopsis* plants. Curr Biol 17: 818–823.

Fang, Y. and D.L. Spector. 2010. Live cell imaging of plants. Cold Spring Harb Protoc 2010: p68.

Fantz, D.A., W.R. Hatfield, G. Horvath, M.K. Kistler, and W.S. Kistler. 2001. Mice with a targeted disruption of the H1t gene are fertile and undergo normal changes in structural chromosomal proteins during spermiogenesis. Biol Reprod 64: 425–431.

Feng, S., S.J. Cokus, V. Schubert, J. Zhai, M. Pellegrini, and S.E. Jacobsen. 2014. Genome-wide Hi-C analyses in wild-type and mutants reveal high-resolution chromatin interactions in *Arabidopsis*. Mol Cell 55: 694–707.

Feng, Z., B. Zhang, W. Ding, X. Liu, D.L. Yang, P. Wei, F. Cao, S. Zhu, F. Zhang, and Y. Mao et al. 2013. Efficient genome editing in plants using a CRISPR/Cas system. Cell Res 23: 1229–1232.

Fortin, J.P. and K.D. Hansen. 2015. Reconstructing A/B compartments as revealed by Hi-C using long-range correlations in epigenetic data. Genome Biol 16: 180.

Fransz, P.F., S. Armstrong, J.H. de Jong, L.D. Parnell, C. van Drunen, C. Dean, P. Zabel, T. Bisseling, and G.H. Jones. 2000. Integrated cytogenetic map of chromosome arm 4S of *A. thaliana*: structural organization of heterochromatic knob and centromere region. Cell 100: 367–376.

Gantt, J.S. and T.R. Lenvik. 1991. Arabidopsis thaliana H1 histones. Analysis of two members of a small gene family. Eur J Biochem 202: 1029–1039.

Geeven, G., Y. Zhu, B.J. Kim, B.A. Bartholdy, S.M. Yang, T.S. Macfarlan, W.D. Gifford, S.L. Pfaff, M.J. Verstegen, and H. Pinto et al. 2015. Local compartment changes and regulatory landscape alterations in histone H1-depleted cells. Genome Biol 16: 289.

Godde, J.S. and K. Ura. 2009. Dynamic alterations of linker histone variants during development. Int J Dev Biol 53: 215–224.

Grob, S. M.W. Schmid and U. Grossniklaus. 2014. Hi-C analysis in *Arabidopsis* identifies the KNOT, a structure with similarities to the flamenco locus of *Drosophila*. Mol Cell 55: 678–693.

Grob, S., M.W. Schmid, N.W. Luedtke, T. Wicker, and U. Grossniklaus. 2013. Characterization of chromosomal architecture in Arabidopsis by chromosome conformation capture. Genome Biol 14: R129.

Harris, C.J., M. Scheibe, S.P. Wongpalee, W. Liu, E.M. Cornett, R.M. Vaughan, X. Li, W. Chen, Y. Xue, and Z. Zhong et al. 2018. A DNA methylation reader complex that enhances gene transcription. Science 362: 1182–1186.

Herrera, J.E., K.L. West, R.L. Schiltz, Y. Nakatani, and M. Bustin. 2000. Histone H1 is a specific repressor of core histone acetylation in chromatin. Mol Cell Biol 20: 523–529.

Huang, Y., Z. Xu, S. Xiong, G. Qin, F. Sun, J. Yang, T.F. Yuan, L. Zhao, K. Wang, and Y.X. Liang et al. 2018. Dual extra-retinal origins of microglia in the model of retinal microglia repopulation. Cell Discov 4: 9.

Huff, J.T. and D. Zilberman. 2014. Dnmt1-independent CG methylation contributes to nucleosome positioning in diverse eukaryotes. Cell 156: 1286–1297.

Jin, F., Y. Li, J.R. Dixon, S. Selvaraj, Z. Ye, A.Y. Lee, C.A. Yen, A.D. Schmitt, C.A. Espinoza, and B. Ren. 2013. A high-resolution map of the three-dimensional chromatin interactome in human cells. Nature 503: 290–294.

Kornberg, R.D. 1974. Chromatin structure: a repeating unit of histones and DNA. Science 184: 868–871.

Kowalski, A. and J. Palyga. 2012. Linker histone subtypes and their allelic variants. Cell Biol Int 36: 981–996.

Krishnakumar, R., M.J. Gamble, K.M. Frizzell, J.G. Berrocal, M. Kininis, and W.L. Kraus. 2008. Reciprocal binding of PARP-1 and histone H1 at promoters specifies transcriptional outcomes. Science 319: 819–821.

Langmead, B. and S.L. Salzberg. 2012. Fast gapped-read alignment with Bowtie 2. Nat Methods 9: 357–359.

Law, J.A. and S.E. Jacobsen. 2010. Establishing, maintaining and modifying DNA methylation patterns in plants and animals. Nat Rev Genet 11: 204–220.

Lee, C.K., Y. Shibata, B. Rao, B.D. Strahl, and J.D. Lieb. 2004. Evidence for nucleosome depletion at active regulatory regions genome-wide. Nat Genet 36: 900–905.

Lee, J.Y. and T.H. Lee. 2012. Effects of DNA methylation on the structure of nucleosomes. J Am Chem Soc 134: 173–175.

Li, G., S. Liu, J. Wang, J. He, H. Huang, Y. Zhang, and L. Xu. 2014. ISWI proteins participate in the genome-wide nucleosome distribution in Arabidopsis. Plant J 78: 706–714.

Lieberman-Aiden, E., N.L. van Berkum, L. Williams, M. Imakaev, T. Ragoczy, A. Telling, I. Amit, B.R. Lajoie, P.J. Sabo, and M.O. Dorschner et al. 2009. Comprehensive mapping of long-range interactions reveals folding principles of the human genome. Science 326: 289–293.

Lin, D., P. Hong, S. Zhang, W. Xu, M. Jamal, K. Yan, Y. Lei, L. Li, Y. Ruan, and Z.F. Fu et al. 2018. Digestion-ligation-only Hi-C is an efficient and cost-effective method for chromosome conformation capture. Nat Genet 50: 754–763.

Lin, Q. A. Sirotkin and A.I. Skoultchi. 2000. Normal spermatogenesis in mice lacking the testis-specific linker histone H1t. Mol Cell Biol 20: 2122–2128.

Lister, R., R.C. O’Malley, J. Tonti-Filippini, B.D. Gregory, C.C. Berry, A.H. Millar, and J.R. Ecker. 2008. Highly integrated single-base resolution maps of the epigenome in *Arabidopsis*. Cell 133: 523–536.

Liu, C., C. Wang, G. Wang, C. Becker, M. Zaidem, and D. Weigel. 2016. Genome-wide analysis of chromatin packing in *Arabidopsis thaliana* at single-gene resolution. Genome Res 26: 1057–1068.

Liu, C., Y.J. Cheng, J.W. Wang, and D. Weigel. 2017. Prominent topologically associated domains differentiate global chromatin packing in rice from Arabidopsis. Nat Plants 3: 742–748.

Love, M.I. W. Huber and S. Anders. 2014. Moderated estimation of fold change and dispersion for RNA-seq data with DESeq2. Genome Biol 15: 550.

Luger, K., A.W. Mader, R.K. Richmond, D.F. Sargent, and T.J. Richmond. 1997. Crystal structure of the nucleosome core particle at 2.8 A resolution. Nature 389: 251–260.

Lyons, D.B. and D. Zilberman. 2017. DDM1 and Lsh remodelers allow methylation of DNA wrapped in nucleosomes. Elife 6.

Maclean, J.A., A. Bettegowda, B.J. Kim, C.H. Lou, S.M. Yang, A. Bhardwaj, S. Shanker, Z. Hu, Y. Fan, and S. Eckardt et al. 2011. The rhox homeobox gene cluster is imprinted and selectively targeted for regulation by histone h1 and DNA methylation. Mol Cell Biol 31: 1275–1287.

Meaburn, K.J. and T. Misteli. 2007. Cell biology: chromosome territories. Nature 445: 379–781.

Mukherjee, A. and R.N. Mukherjea. 1988. Kinetic regulation of hexokinase activity in a heterogeneous branched bienzyme system. Biochim Biophys Acta 954: 126–136.

Nora, E.P., B.R. Lajoie, E.G. Schulz, L. Giorgetti, I. Okamoto, N. Servant, T. Piolot, N.L. van Berkum, J. Meisig, and J. Sedat et al. 2012. Spatial partitioning of the regulatory landscape of the X-inactivation centre. Nature 485: 381–385.

Ozturk, N., I. Singh, A. Mehta, T. Braun, and G. Barreto. 2014. HMGA proteins as modulators of chromatin structure during transcriptional activation. Front Cell Dev Biol 2: 5.

Parnell, T.J. J.T. Huff and B.R. Cairns. 2008. RSC regulates nucleosome positioning at Pol II genes and density at Pol III genes. EMBO J 27: 100–110.

Phillips, J.E. and V.G. Corces. 2009. CTCF: master weaver of the genome. Cell 137: 1194–1211.

Pontvianne, F., M.C. Carpentier, N. Durut, V. Pavlistova, K. Jaske, S. Schorova, H. Parrinello, M. Rohmer, C.S. Pikaard, and M. Fojtova et al. 2016. Identification of Nucleolus-Associated Chromatin Domains Reveals a Role for the Nucleolus in 3D Organization of the A. thaliana Genome. Cell Rep 16: 1574–1587.

Przewloka, M.R., A.T. Wierzbicki, J. Slusarczyk, M. Kuras, K.D. Grasser, C. Stemmer, and A. Jerzmanowski. 2002. The “drought-inducible” histone H1s of tobacco play no role in male sterility linked to alterations in H1 variants. Planta 215: 371–379.

Rabanal, F.A., T. Mandakova, L.M. Soto-Jimenez, R. Greenhalgh, D.L. Parrott, S. Lutzmayer, J.G. Steffen, V. Nizhynska, R. Mott, and M.A. Lysak et al. 2017. Epistatic and allelic interactions control expression of ribosomal RNA gene clusters in Arabidopsis thaliana. Genome Biol 18: 75.

Rao, S.S., M.H. Huntley, N.C. Durand, E.K. Stamenova, I.D. Bochkov, J.T. Robinson, A.L. Sanborn, I. Machol, A.D. Omer, and E.S. Lander et al. 2014. A 3D map of the human genome at kilobase resolution reveals principles of chromatin looping. Cell 159: 1665–1680.

Rea, M., W. Zheng, M. Chen, C. Braud, D. Bhangu, T.N. Rognan, and W. Xiao. 2012. Histone H1 affects gene imprinting and DNA methylation in Arabidopsis. Plant J 71: 776–786.

Russanova, V.R. C.T. Driscoll and B.H. Howard. 1995. Adenovirus type 2 preferentially stimulates polymerase III transcription of Alu elements by relieving repression: a potential role for chromatin. Mol Cell Biol 15: 4282–4290.

Rutowicz, K., M. Lirski, B. Mermaz, G. Teano, J. Schubert, I. Mestiri, M.A. Kroten, T.N. Fabrice, S. Fritz, and S. Grob et al. 2019. Linker histones are fine-scale chromatin architects modulating developmental decisions in Arabidopsis. Genome Biol 20: 157.

Rutowicz, K., M. Puzio, J. Halibart-Puzio, M. Lirski, M. Kotlinski, M.A. Kroten, L. Knizewski, B. Lange, A. Muszewska, and K. Sniegowska-Swierk et al. 2015. A Specialized Histone H1 Variant Is Required for Adaptive Responses to Complex Abiotic Stress and Related DNA Methylation in *Arabidopsis*. Plant Physiol 169: 2080–2101.

Sala, A., M. Toto, L. Pinello, A. Gabriele, V. Di Benedetto, A.M. Ingrassia, B.G. Lo, V. Di Gesu, R. Giancarlo, and D.F. Corona. 2011. Genome-wide characterization of chromatin binding and nucleosome spacing activity of the nucleosome remodelling ATPase ISWI. EMBO J 30: 1766–1777.

Servant, N., N. Varoquaux, B.R. Lajoie, E. Viara, C.J. Chen, J.P. Vert, E. Heard, J. Dekker, and E. Barillot. 2015. HiC-Pro: an optimized and flexible pipeline for Hi-C data processing. Genome Biol 16: 259.

Seymour, M., L. Ji, A.M. Santos, M. Kamei, T. Sasaki, E.Y. Basenko, R.J. Schmitz, X. Zhang, and Z.A. Lewis. 2016. Histone H1 Limits DNA Methylation in *Neurospora crassa*. G3 (Bethesda) 6: 1879–1889.

Shi, L., J. Wang, F. Hong, D.L. Spector, and Y. Fang. 2011. Four amino acids guide the assembly or disassembly of *Arabidopsis* histone H3.3-containing nucleosomes. Proc Natl Acad Sci U S A 108: 10574–10578.

Sirotkin, A.M., W. Edelmann, G. Cheng, A. Klein-Szanto, R. Kucherlapati, and A.I. Skoultchi. 1995. Mice develop normally without the H1(0) linker histone. Proc Natl Acad Sci U S A 92: 6434–6438.

Song, F., P. Chen, D. Sun, M. Wang, L. Dong, D. Liang, R.M. Xu, P. Zhu, and G. Li. 2014. Cryo-EM study of the chromatin fiber reveals a double helix twisted by tetranucleosomal units. Science 344: 376–380.

Sun, L., Y. Jing, X. Liu, Q. Li, Z. Xue, Z. Cheng, D. Wang, H. He, and W. Qian. 2020. Heat stress-induced transposon activation correlates with 3D chromatin organization rearrangement in *Arabidopsis*. Nat Commun 11: 1886.

Thoma, F. T. Koller and A. Klug. 1979. Involvement of histone H1 in the organization of the nucleosome and of the salt-dependent superstructures of chromatin. J Cell Biol 83: 403–427.

Wang, C., C. Liu, D. Roqueiro, D. Grimm, R. Schwab, C. Becker, C. Lanz, and D. Weigel. 2015. Genome-wide analysis of local chromatin packing in Arabidopsis thaliana. Genome Res 25: 246–256.

Wei, T. and M.A. O’Connell. 1996. Structure and characterization of a putative drought-inducible H1 histone gene. Plant Mol Biol 30: 255–268.

Weiner, A., A. Hughes, M. Yassour, O.J. Rando, and N. Friedman. 2010. High-resolution nucleosome mapping reveals transcription-dependent promoter packaging. Genome Res 20: 90–100.

Wierzbicki, A.T. and A. Jerzmanowski. 2005. Suppression of histone H1 genes in *Arabidopsis* results in heritable developmental defects and stochastic changes in DNA methylation. Genetics 169: 997–1008.

Willcockson, M.A., S.E. Healton, C.N. Weiss, B.A. Bartholdy, Y. Botbol, L.N. Mishra, D.S. Sidhwani, T.J. Wilson, H.B. Pinto, and M.I. Maron et al. 2020. H1 histones control the epigenetic landscape by local chromatin compaction. Nature 589: 293–298.

Wollmann, H., H. Stroud, R. Yelagandula, Y. Tarutani, D. Jiang, L. Jing, B. Jamge, H. Takeuchi, S. Holec, and X. Nie et al. 2017. The histone H3 variant H3.3 regulates gene body DNA methylation in Arabidopsis thaliana. Genome Biol 18: 94.

Yan, L., S. Wei, Y. Wu, R. Hu, H. Li, W. Yang, and Q. Xie. 2015. High-Efficiency Genome Editing in Arabidopsis Using YAO Promoter-Driven CRISPR/Cas9 System. Mol Plant 8: 1820–1823.

Yang, S.M., B.J. Kim, T.L. Norwood, and A.I. Skoultchi. 2013. H1 linker histone promotes epigenetic silencing by regulating both DNA methylation and histone H3 methylation. Proc Natl Acad Sci U S A 110: 1708–1713.

Yuan, G.C., Y.J. Liu, M.F. Dion, M.D. Slack, L.F. Wu, S.J. Altschuler, and O.J. Rando. 2005. Genome-scale identification of nucleosome positions in S. cerevisiae. Science 309: 626–630.

Yusufova, N., A. Kloetgen, M. Teater, A. Osunsade, J.M. Camarillo, C.R. Chin, A.S. Doane, B.J. Venters, S. Portillo-Ledesma, and J. Conway et al. 2020. Histone H1 loss drives lymphoma by disrupting 3D chromatin architecture. Nature 589: 299–305.

Yusufova, N., A. Kloetgen, M. Teater, A. Osunsade, J.M. Camarillo, C.R. Chin, A.S. Doane, B.J. Zemach, A., I.E. McDaniel, P. Silva, and D. Zilberman. 2010. Genome-wide evolutionary analysis of eukaryotic DNA methylation. Science 328: 916–919.

Zemach, A., M.Y. Kim, P.H. Hsieh, D. Coleman-Derr, L. Eshed-Williams, K. Thao, S.L. Harmer, and D. Zilberman. 2013. The *Arabidopsis* nucleosome remodeler DDM1 allows DNA methyltransferases to access H1-containing heterochromatin. Cell 153: 193–205.

Zhang, H., R. Zheng, Y. Wang, Y. Zhang, P. Hong, Y. Fang, G. Li, and Y. Fang. 2019. The effects of *Arabidopsis* genome duplication on the chromatin organization and transcriptional regulation. Nucleic Acids Res 47: 7857–7869.

Zhang, L., W.J. Xie, S. Liu, L. Meng, C. Gu, and Y.Q. Gao. 2017. DNA Methylation Landscape Reflects the Spatial Organization of Chromatin in Different Cells. Biophys J 113: 1395–1404.

